# Rod-cone signal interference impairs mesopic motion discriminability in a model circuit

**DOI:** 10.64898/2025.12.26.696616

**Authors:** Adree Songco Aguas, Fred Rieke, Gabrielle J. Gutierrez

## Abstract

Under mesopic conditions, such as dawn or dusk, signals from both rod and cone photoreceptors contribute to perception. These parallel inputs are combined within the retina before being sent to subsequent visual areas. The integration of these kinetically-distinct parallel signals poses unique challenges for human vision. Though previous behavioral studies have found that dim lighting conditions specifically impair motion perception in human subjects, the origin of this dependence is unclear. In the present study, we create a model circuit that predicts ganglion cell responses to moving stimuli by incorporating electrophysiologically-derived circuit components into a Hassenstein-Reichardt correlator, a classical motion-detection model. The model circuit demonstrates that interactions between rod- and cone-derived signals negatively impact the encoding of a moving object’s direction under mesopic conditions. Furthermore, we found that the model circuit could enhance its motion discriminability if it was only sensitive to the cone-activating components of the stimuli. We conclude that rod-cone signal interference occurring at the lowest level of vision has an impact on motion direction discrimination, a higher-level task with relevance for behavior.

## Introduction

In the transition between dim and bright lighting, known as mesopic lighting conditions, rods and cones are simultaneously activated. Mesopic conditions pose a unique challenge to human vision. Because mesopic vision relies on parallel streams of visual input derived from both rod and cone photoreceptors, the retinal circuitry is in a state of transition, balancing the gain of rod-mediated signals that are approaching saturation with the gain of cone-mediated signals that are barely emergent (reviewed by Buck, 2004, Buck, 2014, Grimes et al., 2018, Stockman and Sharpe 2006). Post-photoreceptor circuitry processes the rod and cone inputs in parallel before they are combined to shape retinal outputs (Gouras and Link, 1966, Enroth-Cugell et al., 1977). Consequently, it is not surprising that deficits in mesopic vision are a first symptom in many visual diseases (Petzold and Plant 2006, Arden and Hogg 1985).

Mesopic vision provides a rare opportunity to examine how parallel processing (Kolb and Nelson, 2003; Grimes et al, 2018), a common computational schema in neural circuits, influences human perception. In previous work, electrophysiological recordings have been used to develop a model that predicts retinal ganglion cell responses to rod and cone inputs (Songco-Aguas et al., 2023, Grimes et al., 2018, Grimes et al., 2015). The rod-cone retinal model captures two features of how parallel rod and cone pathways respond to contrived time-varying stimuli such as Gaussian noise and sinusoidal waves: 1) there are distinct kinetics between rod- and cone-derived signals and 2) these parallel signals are combined prior to a shared rectifying or thresholding nonlinearity. These features are consistent with electrophysiological and human psychophysical data showing an unexpected destructive interference of rod- and cone-mediated signals (Grimes et al., 2015, MacLeod, 1972, reviewed by Stockman and Sharpe, 2006).

Models based on physiology are a useful tool for uncovering new mechanistic connections between parallel signal processing and visual perception. Here we focus specifically on motion. Though several human behavioral studies have found that lighting conditions affect the accurate perception of motion (Bilino et al., 2008, Yoshimoto et al., 2013, Yoshimoto et al., 2016, Mayeur et al., 2008, Grossman and Blake, 1999, Sepulveda et al., 2021), the origin of this dependence is unclear. In all of these cases, the impairments primarily affect the perception of translational motion in the periphery. Compared to photopic or scotopic conditions, human observers had particular difficulty in accurately perceiving motion direction for stimuli that mimicked a walking human figure under mesopic conditions (Bilino et al. 2008). Mesopic conditions also uniquely eliminated motion priming in which the perceived direction of motion of an ambiguous stimulus depends on a prior priming motion stimulus (Yoshimoto et al., 2013, Yoshimoto et al., 2016). Here we test the hypothesis that differences in the kinetics of rod- and cone-derived responses in mesopic conditions produce ambiguities in motion discrimination that may contribute to these and related perceptual phenomena (Songco-Aguas et al., 2023).

There are several classical model circuits for motion detection (Barlow and Levick, 1965, Adelson and Movshon, 1982, reviewed in Borst and Egelhaaf, 1989). Fundamentally, these models compute motion direction based on the correlated spatial and temporal changes in brightness that occur when a moving object passes in front of an array of photoreceptors. The Hassenstein-Reichardt correlator consists of two neighboring simulated photoreceptors, or more generally, two neighboring input channels. The circuit correlates the intensity of brightness between the two input channels after the output of one is passed through a temporal low-pass filter that delays its signals relative to the other (Hassenstein and Reichardt, 1956, Reichardt, 1961, Borst and Euler, 2011, Borst and Euler, 2011). It is a relatively simple model that captures the spatio-temporal correlations underlying motion; although it was originally derived from insect optomotor behavior, it is general and abstract enough to model visual motion detection in other species (Borst and Helmstaedter, 2015, Borst and Egelhaaf, 1989, Frechette et al, 2005).

In the context of our study, the Hassenstein-Reichardt model is simply a readout of perceived motion direction of a moving stimulus given rod-cone interactions. We examine how parallel rod and cone signal processing influences the retinal circuit’s ability to accurately ascertain the direction of a moving stimulus. Specifically, by incorporating the rod-cone model circuit into a Hassenstein-Reichardt correlator, we examine the impact of kinetic differences between rod- and cone-derived signals on motion processing in mesopic conditions. Rod-derived signals are slower and have a stronger attenuation of low temporal frequencies than cone-derived signals (see Fig. 1). We first demonstrate the model’s ability to discriminate motion directions across a range of motion speeds for different circuit architectures: circuits that are exclusively responsive to either rod or cone input, and circuits that process both rod and cone mediated motion simultaneously. We hypothesize that the perceptual deficits observed in mesopic conditions are a consequence of the signal interference between rod and cone mediated signals at specific time scales. In comparing the performance differences across these circuits, we gain insight into how the physiologically-derived model retinal circuit degrades motion discrimination ability and potentially contributes to the decreased perceptual acuity observed in human subjects. By testing motion stimuli across a range of motion speeds (i.e. pulse delays), we examine how the known differences in signaling kinetics between rods and cones influence motion processing. Finally, we examine how the salience of motion stimuli impacts motion direction discriminability. Altogether, these findings expand our understanding of how parallel rod-cone processes underlie the perception of stimuli that can affect human behavior.

**Figure 1.**
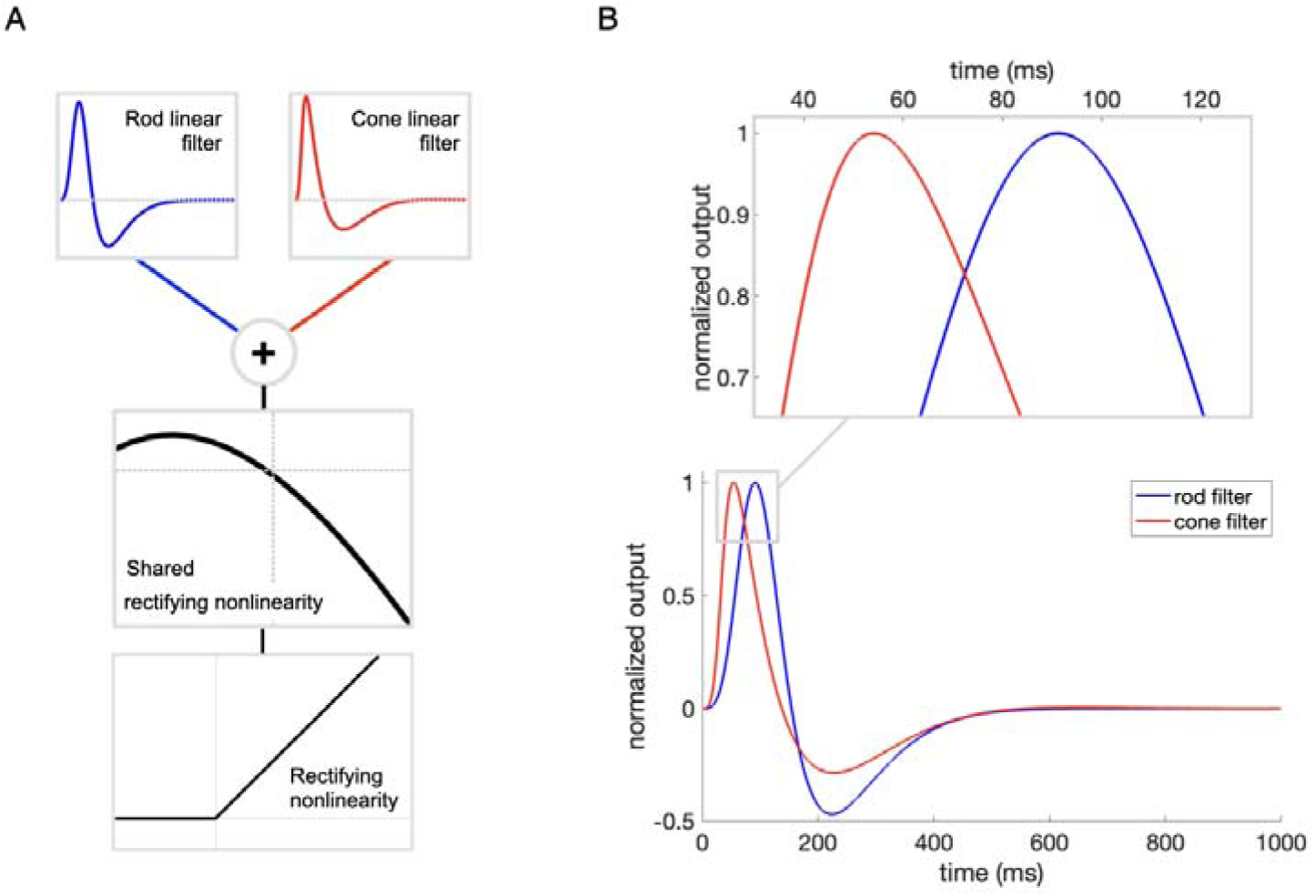
Rod-cone LN circuit model. A: Rod-cone linear-nonlinear cascade model. A two-pronged model of an ON parasol retinal ganglion cell with rod and cone input. The linear filters capture the unique kinetics of rod- and L-cone-derived responses in mesopic conditions (blue: rod; red: cone). After linear filtering, rod and cone signals are summed, then passed to a shared nonlinearity and then a rectifying nonlinearity. The final output of the rod-cone retinal model represents the spiking rate of a retinal ganglion cell. B: Rod and cone filter kinetics. The rod filter (blue) is slower and has an overall more biphasic shape than the cone filter (red). The inset shows that the delay in time-to-peak between rod and cone filters is about 35 ms.

## Methods

### Retinal model

The rod-cone retinal model was derived from electrophysiological recordings conducted on whole mount preparations of isolated non-human primate retina, as previously described (Dunn et al., 2007; Trong and Rieke, 2008). These recordings were taken from ON parasol ganglion cells. The circuit components of the rod-cone linear-nonlinear model (Figure 1) were derived from voltage-clamp recordings of full-field white noise (0-40 Hz bandwidth) using blue LEDs (peak power at 460 nm) and red LEDs (peak power at 640 nm) to isolate rod and L-cone responses, respectively, as previously described (Songco-Aguas et al., 2023, Grimes et al., 2015, Chichilnisky 2001). Circuit components of the model were verified by taking the model’s explained variance from data with the same noise stimuli presented.

The linear-nonlinear model responses to motion stimuli, *s*(*t*), were simulated as follows. First, rod and cone stimuli, *s*_rod_(*t*) and *s*_cone_(*t*), are convolved with their respective linear filters, h_rod_ and h_cone_.

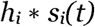

Filtered signals are then summed. The summed signal is passed into a shared nonlinearity that was fit from the ganglion cell responses with a third degree polynomial function (see Fig. 1A, middle panel); the output of this nonlinearity is an estimate of the cell’s excitatory synaptic input. A second nonlinearity, a rectified linear function, *r*, converts synaptic currents into retinal ganglion cell spike rates (see Fig. 1A, bottom panel).

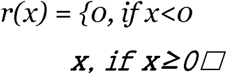

Instantaneous currents below 0 amps do not result in spiking, but above that threshold, the spike rate linearly increases with increasing current.

To define the shared nonlinearity between rod and cone components of the linear-nonlinear model for a single cell (Schwartz et al, 2012; Dunn et al, 2006), the horizontal axis of the measured cone nonlinearity is scaled relative to the measured rod nonlinearity until they overlap, and then the cone filter is scaled accordingly, as previously described (Songco-Aguas et al., 2023).

The rod-cone retinal circuit incorporated into the Hassenstein-Reichardt model (see following section) was fitted with data from five ganglion cells. The linear and nonlinear elements were fit to a respective polynomial function for each recorded retinal ganglion cell, then the parameter fits for these polynomial functions were averaged across cells.

### Hassenstein-Reichardt correlator

The architecture of our Hassenstein-Reichardt model is illustrated in Figure 2 (see Supplementary Fig. S1 for illustration of classical Hassenstein-Reichardt model). The input channels interact with each other with a correlator time delay (Δt). For example, the real-time response of the right input channel (*r_right_*) is multiplied with the time-delayed response of the left input channel ( ). By mirroring this process (multiplying the real-time response of the left input channel, *r_left_*, with the time-delayed response of the right input channel,), then subtracting the products from one another, we remove the parts of the correlator’s response that result from direction-irrelevant correlations in the stimulus (Borst and Egelhaaf 1989). The final output of the Hassenstein-Reichardt model is calculated as:

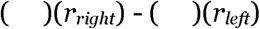

**Figure 2.**
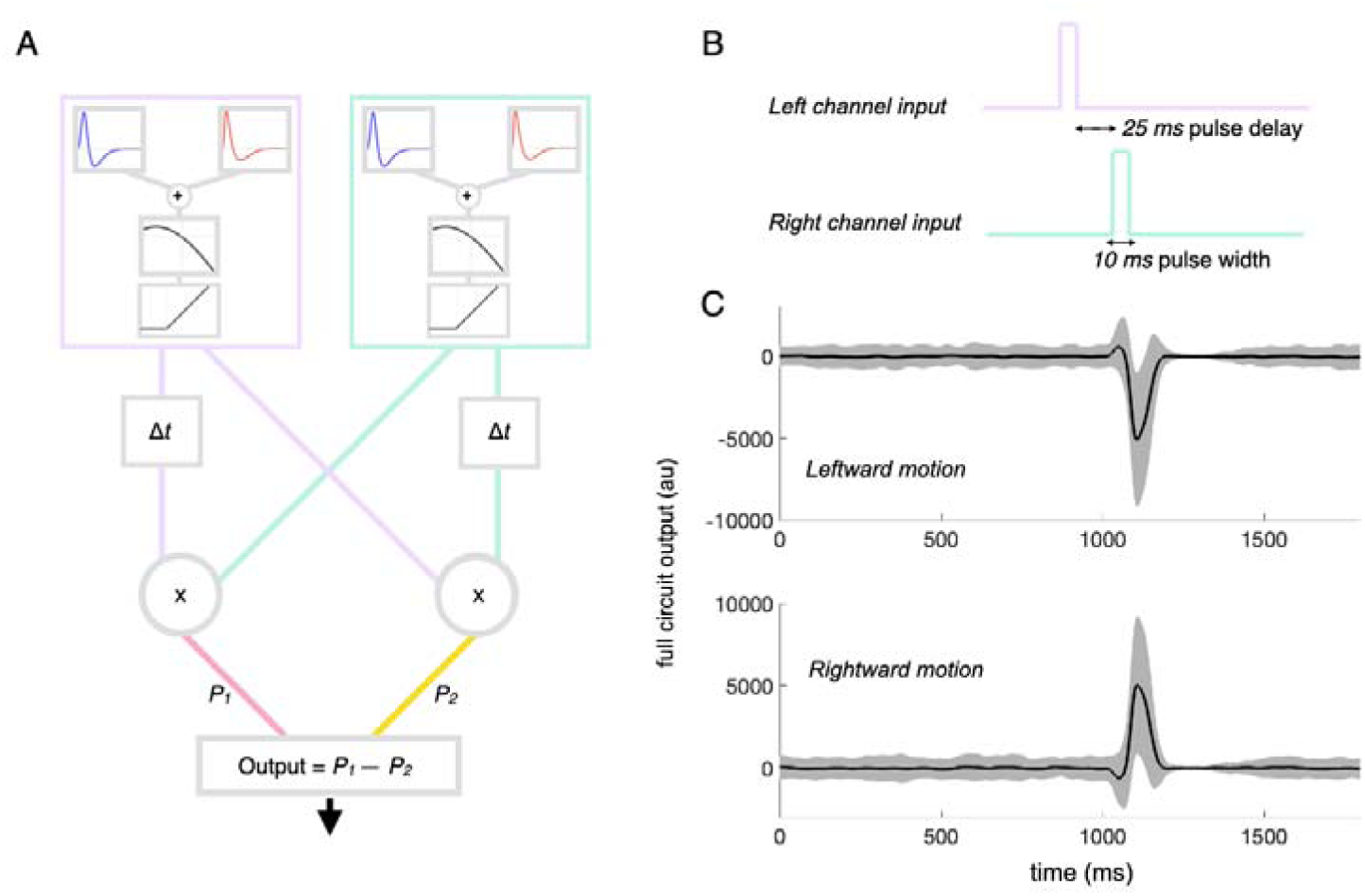
Motion correlator with rod-cone retinal model input channels. A: Hassenstein-Reichardt correlator with the rod-cone retinal circuit model as the input channels computes motion from rod-cone activating stimuli. Stimuli pass through one input channel and then the other after some time delay (Δs) corresponding to motion velocity. Real-time and time-delayed (Δt) signals from opposite input channels are multiplied. For example, *P*_1_ is the product of a time-delayed signal from the left input channel and the real-time signal from the right input channel. The difference between the products (*P*^1^ and *P*_2_) is computed. The output is an expression of the motion of direction. B: Motion stimulus trace for a single trial with a 10 ms pulse width and a 25 ms pulse delay, in which the left input channel is presented with a 10 ms wide stimulus pulse, and then 25 ms later, the right input channel is stimulated with the 10 ms pulse. Noise is added to these pulses before they are processed by the rod-cone circuit. C: The average response traces for leftward (top) and rightward (bottom) motion across 100 trials. The preferred direction of motion elicits a positive peaked output. Gray cloud shows standard deviation of mean at a given time point.

The pulse delay, Δs, is a free parameter that controls the time delay between the arrival of the stimulus at one input channel and its arrival at the other input channel. In other words, the pulse delay corresponds to the motion speed of the stimulus. The **pulse width**, *w*, is another free parameter and it controls the duration of the stimulus (i.e. the width of the stimulus pulse, or the overall angular size of the stimulus). Theoretically, a Hassenstein-Reichardt circuit would be most sensitive to the pulse delays that match the correlator time delay. As the pulse delay (Δ*s*) diverges from the correlator time delay (Δ*t*), wider and wider pulses widths (*w*) are needed for the circuit to register the stimulus as a moving stimulus. This takes the form of a piecewise linear relationship.

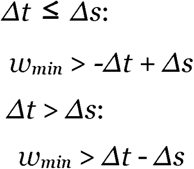

We illustrate this function in Supplementary Figure S1. The piecewise linear relationship describes the Hassenstein-Reichardt correlator without the filter kinetics or nonlinearities, but once those are included, this relationship is no longer guaranteed to be linear due to the nonlinearities inherent in the rod-cone retinal model circuit.

In our simulated experiments, we chose a fixed Hassenstein-Reichardt correlator time delay of 50 ms (see Supplementary Fig. S1), with the exception of the simulated experiments shown in Supplementary Figure S2. For that experiment, we tested circuits with time delays of 25 ms and 75 ms to examine how the time delay affected motion discriminability using our full model circuits. We chose a correlator time delay of 50 ms because this is comparable to the temporal separation between rod and cone filters (see Fig. 1) and hence corresponds to time offsets in which rod and cone signals may interact.

### Stimulus generation and simulated experiments

For each individual trial, a pulse of light was presented to one of the linear-nonlinear model input channels. Then, to simulate spatial distance between two cells, the other input channel received the same pulse of light after a pre-specified pulse delay between 0 ms and 100 ms, corresponding with stimulus velocity (larger pulse delays are slower velocities while smaller pulse delays are faster velocities). To mimic noise in the real responses, Gaussian pink noise with a mean of 0 and a standard deviation of 10% was scaled by the mean intensity of the stimulus and added to the pulse stimulus. Noise was uncorrelated between rod-activating and cone-activating components of the motion stimulus and also uncorrelated between stimuli presented to the left and right input channels. The true direction of motion was based on which input channel received the pulse first.

### Classification and performance

The model output was classified as a leftward or rightward trial with a discriminant vector (Zhao et al, 2024). The discriminant vector for a set of trials with a specific pulse delay was calculated by taking the mean response traces of the model for 100 leftward trials and 100 rightward trials, then subtracting the mean leftward response trace (non-preferred direction) from the mean rightward response trace (preferred direction). This linear discriminant analysis is effective under the conditions of our simulations in which the noise is independent of the stimulus and uncorrelated across responses. This process mirrors the Hassenstein-Reichardt correlator calculation resulting in the preferred direction of motion having a positive polarity.

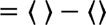

To automatically label motion direction for a single trial, the response trace was projected onto the discriminant vector by taking the dot product. A projection value of zero can be interpreted as a response trace that was perfectly dissimilar from both the mean leftward trace and the mean rightward trace. Projections with a value greater than zero were automatically labeled rightward, while projections with a value less than zero were automatically labeled leftward. We ran 1000 new trials, half with leftward motion and half rightward motion. The model’s ability for motion direction discrimination was measured as the percentage of trials where the automatic labeling captured the true direction of the motion stimulus. Separate probability density functions were fitted to these the projection values of true leftward and true rightward trials to visualize the overlap in trials labeled leftward or rightward. Confidence intervals for the full model output responses were computed using the Clopper-Pearson interval.

### Implementation

All simulations were run in MATLAB version 2024b using a MacBook Pro with an Apple M2 chip and 24 GB of memory. Code is publicly available on the Gutierrez lab’s motion-detection repository, which can be found on https://github.com/Gutierrez-lab/motion-detection.

## Results

### A motion discrimination circuit with rod-cone kinetics

To simulate retinal ganglion cell spike rate traces, we fit a linear-nonlinear cascade model using previously described methods (Songco-Aguas et al., 2023, Grimes et al., 2015, Chichilnisky 2001). Voltage-clamp recordings of ON parasol ganglion cells from non-human primate retina in mesopic conditions were used to fit the linear and nonlinear components of our cascade model. This linear-nonlinear model circuit captures the unique kinetics of rod-derived and cone-derived signals in mesopic lighting conditions. Specifically, the rod filter has a longer time-to-peak than the cone filter–with rod-derived signals delayed by approximately 33 ms relative to cone-derived signals (Fig. 1B). The rod filter is also more biphasic than the cone filter, meaning that the rod response has more of a negative response component after the initial positive response component (Songco-Aguas et al., 2023, Grimes et al., 2018). As a consequence, we would expect that a rod-only linear-nonlinear model compared to a cone-only linear-nonlinear model would be slower in responding to the same stimulus.

In addition to these kinetic filters, the linear-nonlinear model circuit also contains a nonlinearity that shapes ganglion cell inputs. Because the shape of this nonlinearity was found to be consistent between the rod-targeted and cone-targeted electrophysiological recordings, we interpreted this phenomenon to mean that rod and cone signals are integrated prior to this nonlinear stage, such as in the cone bipolar output synapse (Grimes et al., 2015, Songco-Aguas et al., 2018, Fain and Sampath 2018, Gouras and Link 1966). Indeed, this model architecture captures temporal interactions between flashed stimuli that selectively activate rods or cones (Grimes et al., 2015). Models lacking a shared nonlinear component could not capture these interactions. The output of the nonlinearity was converted into retinal ganglion cell spike rates through a rectified linear function. Altogether, we refer to these components of our linear-nonlinear model collectively as the rod-cone retinal model circuit (Fig. 1A; see Methods).

We incorporated our rod-cone retinal models as the two input channels of a Hassenstein-Reichardt correlator (Figure 2A). The Hassenstein-Reichardt correlator detects motion using a “delay-and-compare” computation (Hassenstein and Reichardt, 1956, Reichardt, 1961, Borst and Euler, 2011, Borst and Euler, 2011). It relies on two input channels with identical response properties–in our case two simulated retinal ganglion cells–with a fixed spatial separation between them. This spatial separation is implemented as a temporal delay, Δ*t*. The direction of motion is given by the order in which each input channel receives the stimulus, and the velocity of motion is based on the delay in time between when the first channel and the second channel receive the stimulus, which we refer to as the pulse delay, Δ*s* (Fig. 2B). The output from the left channel is time-delayed then multiplied with the real-time output of the right channel, and vice versa. The resulting polarity in the responses provides the circuit’s directional tuning. In this case, our circuit’s preferred direction of motion is rightward because a rightward stimulus produces a response trace with a positive peak whereas a leftward stimulus produces a negative peak (Fig. 2C). A leftward-preferring circuit would be identical, except at the subtraction stage, which would have P_2_-P_1_ instead (see Figure 2A).

Our motion input consists of a noisy “moving” pulse—that is, a pulse of a given magnitude that activates one of the input channels, then after a period of time, Δ*s*, a second pulse that activates the other input channel. For these initial trials, we chose our pulse magnitude to have a 25% contrast over a fixed mean luminance. These pulses contained both rod-activating and cone-activating stimulus components, and the stimulus was scaled such that both rod and cone wings of the rod-cone retinal model were equal contributors to the spike rate output. Independent Gaussian pink noise was also added to each pulse, unique to each trial.

Noise was uncorrelated between stimuli presented to the left and right input channels, as well as uncorrelated between rod-activating and cone-activating components of the motion stimulus.

Figure 2C shows that the mean output traces of the Hassenstein-Reichardt correlator are opposite in polarity between stimuli moving in its preferred direction (rightward) and its non-preferred direction (leftward) when other stimulus parameters (pulse width and pulse delay) are kept fixed.

We created a systemized method such that the model circuit responses could be decoded and automatically labeled by the “perceived” direction of motion for each trial. To do this, we calculated mean response traces across leftward and rightward trials of a given pulse delay (Fig. 3A; see Methods). We then calculated a discriminant vector by subtracting the mean leftward response trace from the mean rightward response trace. This process mirrors the subtraction stage in the symmetrical Hassenstein-Reichardt model (Fig. 2A) that yields a preferred direction of motion. We then took the output traces across 1000 new trials–half leftward and half rightward–and projected them along the discriminant (Fig. 3A). Automatic direction discrimination was based on whether the projection value of a trial was greater than zero, in which case it was labeled as rightward motion, or whether it was less than zero, in which case it was labeled as leftward motion (Fig. 3B). We computed probability densities of the projection values for the leftward and rightward trials to visualize the overlap between these two distributions (Fig. 3C). This overlap determines what fraction of trials would be erroneously identified (i.e. rightward trials with projection values less than 0 and leftward trials with values greater than 0).

**Figure 3.**
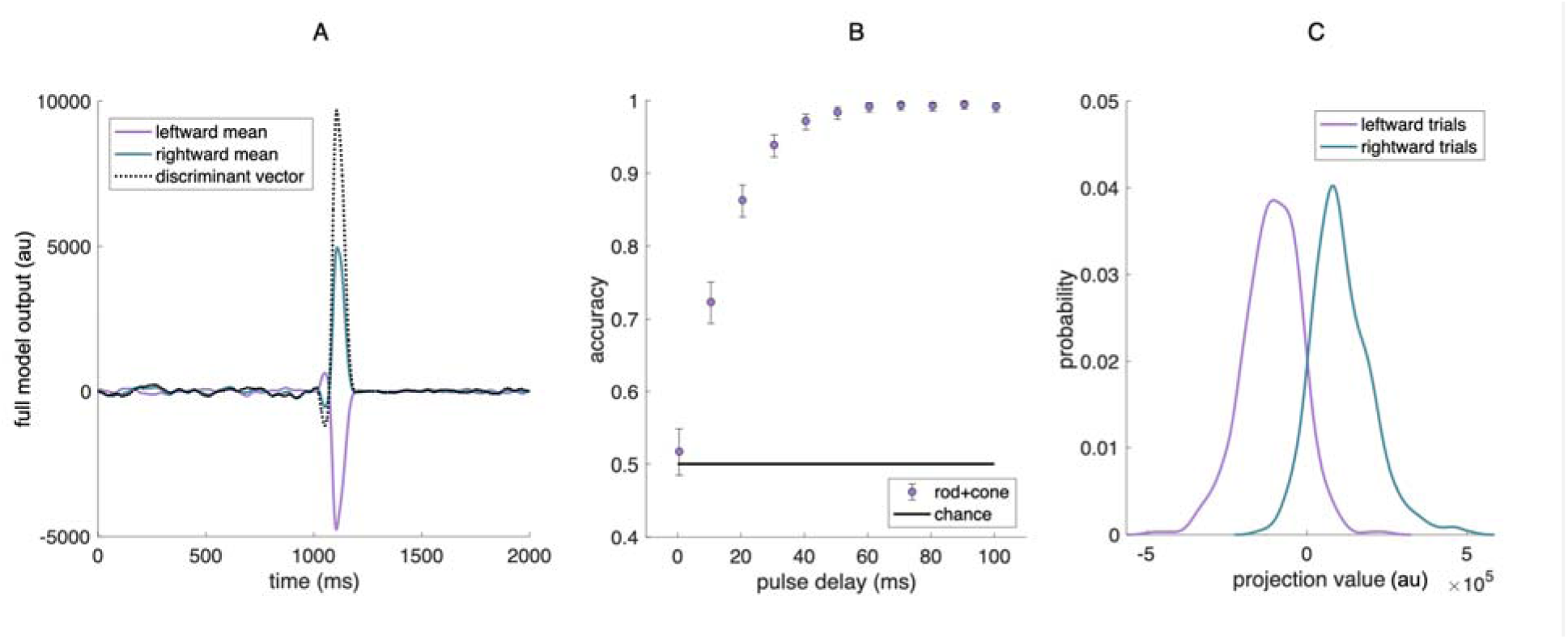
Automatic labeling of motion direction. A: Discriminant vector for a rod-cone stimulus with 25 ms pulse delay and 10ms pulse width (Δ*s* = 25 ms, *w* = 10 ms). The mean output trace of the Hassenstein-Reichardt circuit is computed for both directions of motion and the discriminant vector for the rightward-preferring circuit is the mean rightward motion output minus the mean leftward motion output (see Methods). B: Motion discrimination performance across a range of pulse delays. Accuracy for discriminating motion direction can vary between 0 and 1, where 0 is no accuracy, 1 is complete accuracy, and 0.5 is chance performance (indicated by solid line). Error bars represent the confidence interval with a minimum coverage of 95% of the data, the Clopper-Pearson interval. C: Probability distribution of projections for trials with a 25 ms pulse delay stimulus. Two distinguishable probability distributions arise from fitting the projection values from leftward (purple) and rightward (teal) trials. The overlap between the two distributions represents trials in which leftward and rightward motion are potentially misclassified.

With our automated labeling method, we calculated the circuit’s motion discrimination accuracy from the set of 1000 trials: the fraction of trials in which the model’s automatic label corresponded to the true direction of the presented stimuli (Fig. 3B). As to be expected, the model performs at chance in the absence of a pulse delay (Δ*s* = 0 ms) because the stimulus inputs are presented simultaneously to the left and right input channels; hence, for these trials, there was no actual motion in the stimulus. For pulse delays of 20 ms, the model performed with ∼85% accuracy (i.e., it incorrectly labeled the direction of motion in ∼15% of the trials). Performance plateaued for delays of 60-100ms, while smaller delays resulted in the circuit mislabeling a larger fraction of trials. The model’s inaccuracy with labeling stimuli with a smaller pulse delay (i.e., faster moving stimuli) seemed to correspond to the difference in the time-to-peak of the rod and cone linear filters. Our previous work has demonstrated how responses to time-varying stimuli at specific temporal frequencies can be attenuated due to interference between rod and cone signals (Songco-Aguas et al., 2023). In the next section, we further investigate the effect of rod-cone interference on motion direction discrimination.

### Disentangling the perceptual contributions of rods and cones

Does interference between parallel rod- and cone-derived signals hinder motion discrimination? We answered this question by comparing the performance of the rod-cone circuit with the performance of a circuit in which the input channels were exclusively sensitive to the rod-activating component of the presented stimuli (rod-only) and a circuit in which the input channels were exclusively sensitive to the cone-activating component (cone-only). In effect, these trials were simulations of the peripheral retina with either a rod or cone knockout. We repeated the same range of pulse delays (ie., 0-100ms) as in our previous test. The results are plotted in Figure 4.

**Figure 4.**
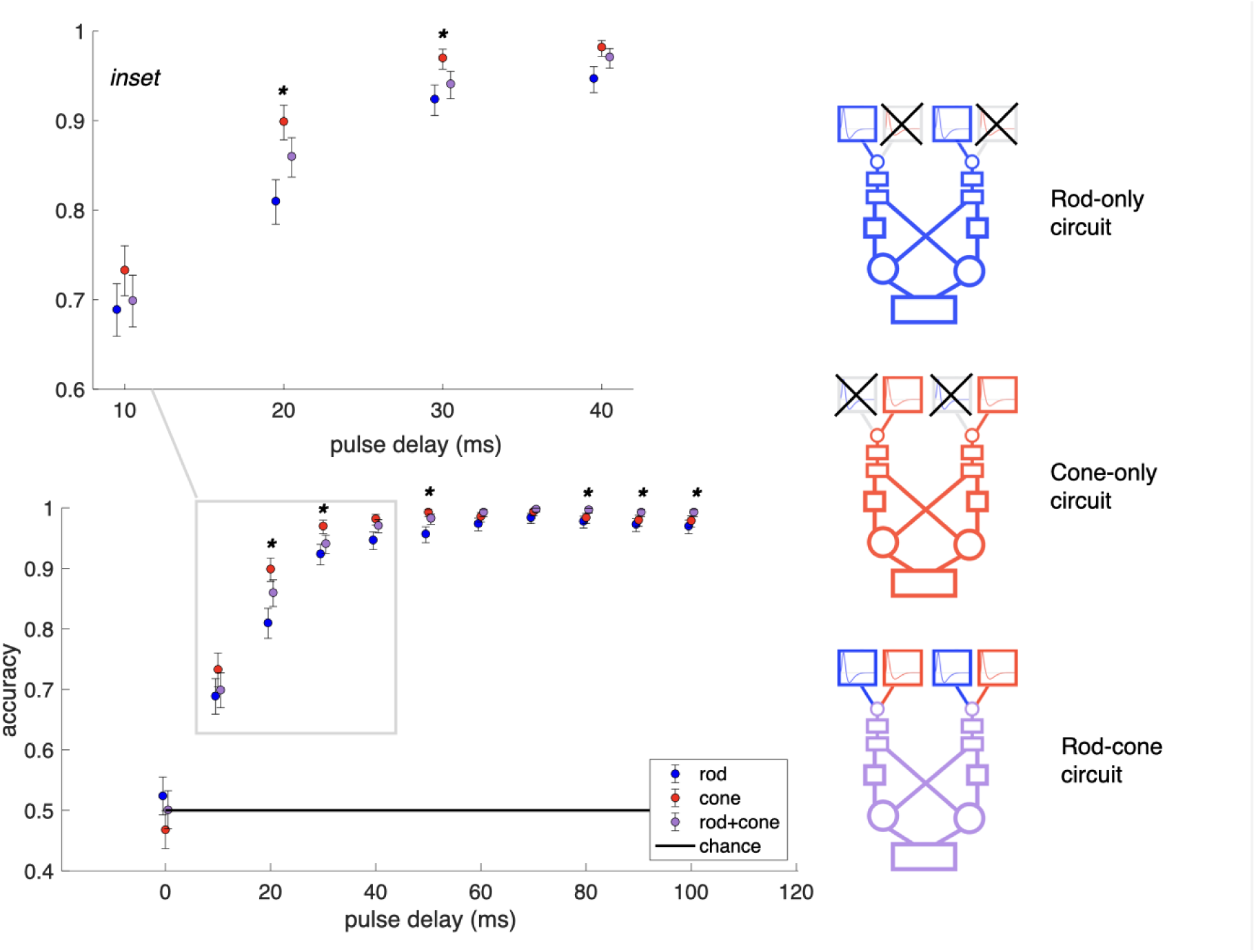
Motion discriminability between simulated knockout models. We compare the discriminability between three different retinal model circuits as input channels to the Hassenstein-Reichardt model, each with a fixed correlator time delay of 50ms (right). The rod-only circuit (blue) is exclusively responsive to the rod-activating components of motion stimuli, the cone-only circuit (red) is exclusively responsive to the cone-activating components, and the rod-cone circuit (purple) is responsive to both. Identical motion stimuli are passed into these unique circuits, and discriminability is computed for each circuit and condition. Inset: for motion stimuli between 20-30 ms, the cone-only circuit outperforms both the combined and rod-only circuits in motion discriminability. For stimuli <10 ms and ≥ 80 ms, performances between the circuits are comparable. Error bars represent the confidence interval with a minimum coverage of 95% of the data (Clopper-Pearson interval). Solid black line indicates chance (a motion discrimination accuracy of 0.5, ie., 50%). Asterisks denote where statistical significance was reached in the comparison between the cone-only circuit performance and the combined rod-cone circuit performance (p ≤ 0.05).

For pulse delays of 20 and 30 ms, we found a separation in motion discrimination performance across the three circuits in which the cone-only circuit outperformed both the rod-cone model circuit and the rod-only circuit (Figure 4 inset). It was unsurprising that the cone-only circuit outperformed the rod-only circuit, since the cone-derived responses are both faster and more monophasic in their kinetics than the rod-derived responses. Ultimately, a faster response time would be better at encoding faster moving stimuli. Additionally, due to the more monophasic shape of the cone filter relative to the rod filter, more of the temporal structure of a given stimulus would be preserved in the output of the circuit.

We had hypothesized that the rod- and cone-derived signals would interfere with motion discriminability not only because of their relative delay in kinetics but because of how they are integrated in the retina. The observation that the cone-only circuit outperformed both rod-cone and rod-only model circuits is in line with this initial hypothesis; that rod-cone signal interactions would impact motion computation. However, the rod-cone circuit had better performance than the rod-only circuit which makes it difficult to determine the extent to which impaired motion discriminability may not only be due to the characteristics of rod response kinetics, but to the offset in time-to-peak response kinetics for rods and cones. We tested a circuit with two cone inputs per channel where one cone filter is shifted relative to the other cone filter by the same amount as the rod filter would have been (∼33 ms). This circuit performed as poorly as the rod-only circuit despite not having any of the broad and biphasic features of the rod response – only the offset in time-to-peak. Thus, the offset between the rod and cone response kinetics, and not merely the broad and biphasic kinetics of the rod response, interferes with motion discrimination.

Our choices for the model parameter values can tune the preferences of the model. A preference for direction of motion is built into the Hassenstein-Reichardt correlator circuit during the subtraction stage (Fig. 2A). At the same time, the choice of correlator time delay can tune the circuit to prefer a range of pulse delays, which correspond to motion velocity, to a certain extent (Supplementary Fig. S2, additional detail in Methods section “Hassenstein-Reichardt correlator”). For pulse delays that exactly match the correlator time delay, the circuit can accurately discriminate motion direction for the largest range of pulse widths. In other words, the model has optimal motion direction discrimination for moving stimuli over a wider range of stimulus sizes when the delay between two pulses, corresponding to motion velocity, matches the Hassenstein-Reichardt correlator’s tuning. For stimuli within the range of preferred pulse delays, the cone-only circuit performance benefits the most. This is again because of the faster-to-peak, more monophasic shape of the cone filter, compared to the rod filter. The cone filter more closely resembles an impulse response than the rod filter, thus encoding the stimulus more faithfully. Conversely, the rod-only circuit consistently performs with the least accuracy because the biphasic shape of the rod responses across the left and right channels reduces the signal, making it more ambiguous. Taken together, these results demonstrate that rod-cone signal interference interacts with both the directionally-tuned and the velocity-tuned components of the motion correlator circuit.

### Stimulus salience and motion discriminability

Which properties of the motion stimulus itself had an impact on the separation in performance between the three different circuits? The size and relative contrast of a stimulus define its salience. A wide stimulus pulse width represents a larger, and thus more salient, stimulus than one with a narrow pulse width. Likewise, a high contrast stimulus is more salient than a low contrast stimulus. As shown in Figure 5, we varied the size and contrast of the motion pulse while keeping all other stimulus parameters fixed. We found that motion direction was more accurately classified by all three variations of the circuit (rod-cone, rod-only, and cone-only) for stimuli with longer pulse widths and greater contrast (Fig. 5, top right panel) . In other words, the model’s discrimination performance generally increased with stimulus salience.

**Figure 5.**
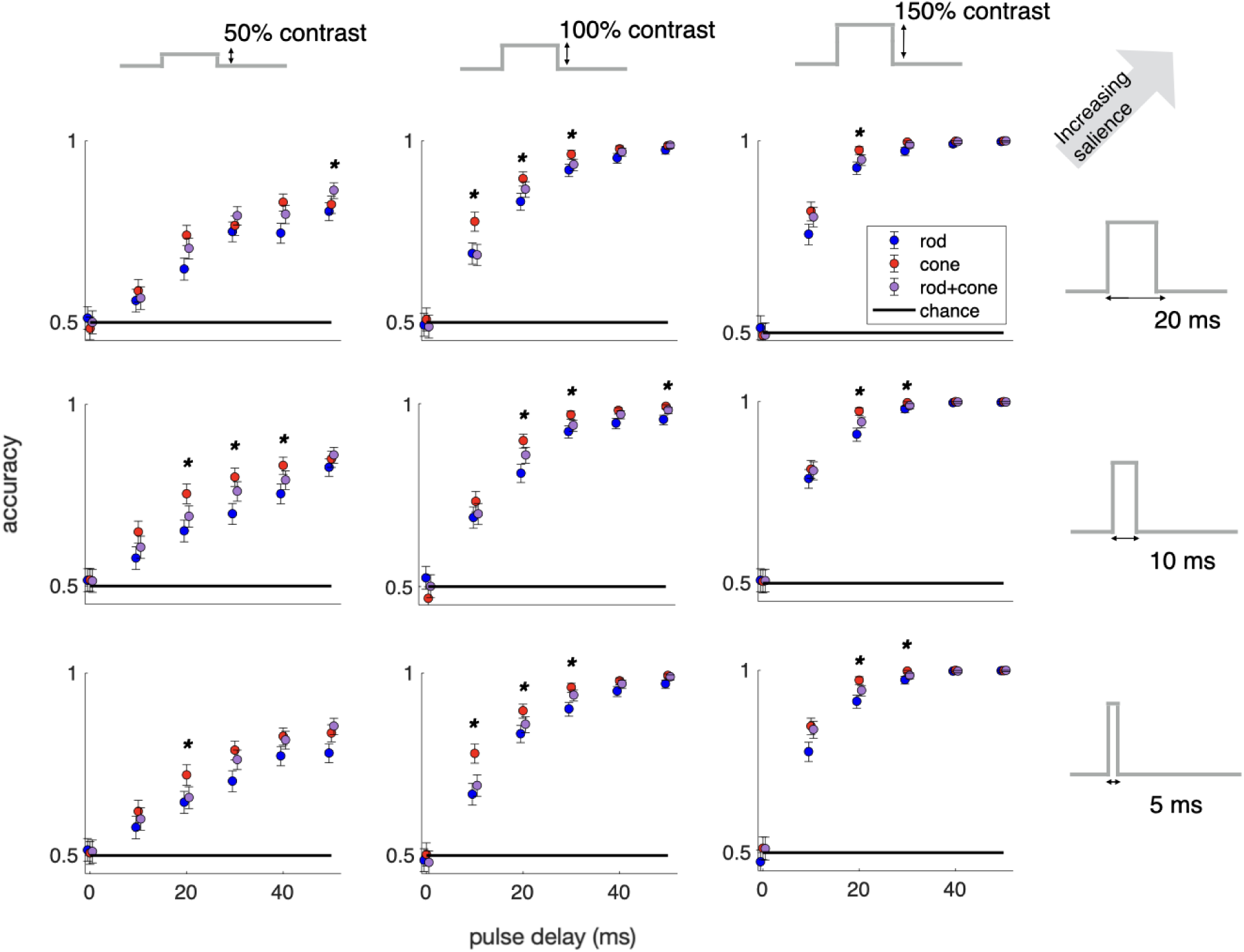
Motion discriminability and stimulus salience. In general, discriminability performance improves with increased salience–specifically with increased stimulus pulse width (vertical) or increased contrast (horizontal). Error bars represent the confidence interval with a minimum coverage of 95% (Clopper-Pearson interval). Solid black line indicates chance (a motion discrimination accuracy of 0.5, ie., 50%). Asterisks denote statistical significance was reached in the comparison between the cone-only circuit performance and the combined rod-cone circuit performance (*p* ≤ 0.05).

Performance suffered for all circuits presented with short pulse widths and low contrast stimuli (Fig. 5, bottom left panel). The stimuli with 150% contrast had an increased range of pulse delays in which all circuits were 100% accurate at labeling motion direction (Fig. 5, right column), compared to the 50% or 100% contrast stimuli. On the other hand, varying the pulse widths had a less pronounced impact on the circuit’s overall performance within a given stimulus contrast.

For most of these conditions, there was still a separation of accuracy between the cone-only circuit relative to rod-only and rod-cone circuits across 20-30 ms pulse delays, that was previously observed for 10 ms pulse widths at 100% contrast. For the 20 ms pulse widths at 50% contrast, however, there was no significant separation in the performances between the models at pulse delays between 20-30 ms.

Ultimately, our model demonstrates that interactions between rod- and cone-derived signals negatively impact the computation of motion direction. When we pull apart rod-cone signal interactions by creating a circuit that is only responsive to cone inputs (ie., essentially a rod knockout circuit), we find that the circuit is able to discriminate motion direction with greater accuracy. For stimulus conditions which further decreased the salience of the motion stimulus, we observe a greater overall degradation in motion discriminability relative to discriminability for more salient stimuli.

## Discussion and Conclusions

Unlike photopic lighting conditions where cone photoreceptors dominate the visual response and scotopic lighting where rod photoreceptors dominate the visual response, mesopic lighting activates both rods and cones (reviewed by (Buck, 2004, Buck, 2014, Grimes et al., 2018)). We hypothesized that destructive interference between rod- and cone-mediated signals in the retina would hinder motion perception given that it hinders the ability to perceive stationary flickering stimuli of certain frequencies (Songco-Aguas et al, 2023). Alternatively, it is possible that rod and cone signals could constructively contribute to motion discrimination by averaging out the independent noise inherent to each rod and cone input pathway, thus improving the signal to noise ratio. To examine this, we incorporated a rod-cone retinal circuit model trained on electrophysiology data into a directionally-sensitive motion-detection circuit. We found that motion discriminability was decreased by the integration of rod- and cone-derived responses for a specific range of pulse delays–namely 20-30 ms–corresponding with the relative delay in kinetics between rod- and cone-derived responses. Less salient stimuli led to greater losses in motion discriminability.

### Simulating motion correlation as a post-retinal processing stage

Our goal was to investigate how interactions between parallel rod- and cone-derived signals impacted the fidelity of retinal outputs, with motion discrimination as an important example. We used a classical motion correlator, the Hassenstein-Reichardt model as a readout of motion. It was originally developed as a model of motion encoding in the insect visual system, and it relies on the correlations between two stimulus inputs to compute the presence of motion for a given stimulus (Reichardt 1961). The classical Hassenstein-Reichardt model has been shown to be a useful model for motion perception generally, including for applications in computer vision (Qiu et al., 2025, Birkoben et al., 2023, Yan et al. 2022, Basch et al. 2010). For our purposes, it serves as a convenient readout to estimate the encoded motion information from two channels of the retinal model. To examine the impact of early retinal processing on motion encoding, we enhanced the Hassenstein-Reichardt model by integrating a simulated rod-cone circuit into the channels. The rod and cone response kinetics and nonlinearities were fit using electrophysiology data collected from non-human primate retina. By including rod and cone linear kinetics and shared nonlinear processing, we explicitly study how signal interactions among those components impact a canonical motion computation.

While the response filters for the rods and cones in our model are physiologically realistic, the correlator time-delay and multiplication components do not have a corresponding physiological interpretation in the primate retina nor further downstream. However, the full model is an abstract emulation of the directionally-tuned and velocity-tuned visual processing that occurs in the visual system, downstream of the retina. The directional tuning is based on the subtraction stage prior to the final output of the Hassenstein-Reichardt model. The velocity tuning comes from the value of the specific correlator time delay. This kind of directional and velocity tuning would be part of post-retinal computations, such as those that occur in the middle temporal (MT) visual area (Maunsell and van Essen, 1983).

Other studies have observed perceptual deficits in motion processing that are attributed to post-retinal processing. For example, subjects have a difficult time reporting the location of a moving object at high speeds, perhaps because of a deficit in the high-level binding of the position of the object and the cue to report the position (Linares et al, 2009). Signals traveling through the parvocellular pathway are slower than those through the magnocellular pathway (Maunsell et al, 1999), allowing for the possible interpretation of the correlator time delay in the Hassenstein-Reichardt model in terms of this post-retinal processing feature; however, our study exclusively concerns motion discrimination, which is generally handled by the magnocellular pathway (Merigan et al., 1991). Our study probed the role of specific retinal interactions in perceptual deficits related to motion discrimination, but future studies may expand on our model to investigate how post-retinal processing interacts with rod-cone interactions.

### Rod-cone signal interference degrades motion discriminability

In our present study, we found that motion processing is degraded by destructive signal interference, as evidenced by the worse performance of the rod-cone model circuit compared to its cone-only counterpart. Poor motion discriminability was to be expected from the rod-only circuit because the slow kinetics of the rod input degrade the fine temporal structure required for the accurate encoding of the time-varying stimulus (Songco-Aguas 2023). Similar to the classic visual stimulus of a flickering spot (MacLeod 1977, Stockman and Sharpe 2006, Songco-Aguas et al., 2023), movement or change of any kind requires the stimulus to vary in time. The lag between rod-mediated responses and cone-mediated responses and subsequent destructive interference ultimately blurs the final temporal structure of the stimulus, preventing accurate stimulus encoding within a channel. In contrast to a flickering spot, our moving stimulus also varied in space. The loss of fidelity in encoding within a channel propagates through the circuit, where a comparison across both channels takes place with a built-in time delay.

In our previous research, we paired electrophysiology with human psychophysics to investigate how the kinetic difference between rod- and cone-mediated signals influences visual perception in certain conditions (Grimes et al., 2015, Songco-Aguas et al., 2023). In one of these studies, we examined how interference between rod- and cone-mediated signaling led to the inability to perceive flickering stimuli in the peripheral retina (Songco-Aguas et al., 2023). We found evidence for destructive interference between these two signal pathways and that we could affect the interference by altering either the temporal frequency of the flickering stimulus or the relative phase delay between the rod-activating and cone-activating components of the stimulus (Songco-Aguas et al., 2023).

That said, our motion-discrimination circuit is not only dependent on the responses from a single channel – it also needs to compare the signals from two channels receiving stimulus input with a delay that represents motion across the visual field. Our interpretation for our model circuit’s performance drop in discriminability is that 1) rod-cone signal interference happens to some extent within a channel, degrading the overall output of single channels, and that 2) this signal degradation is exacerbated by the motion correlator circuit in its multiplication stage. This is akin to how the linear stages of the rod-cone retinal model initially cause the interference, while the consequent nonlinear stages heighten the effect of the signal interference downstream. Specifically, the multiplication stage of the correlator is a nonlinear process that emphasizes correlations in a time-varying motion stimulus (Dror, O’Carroll, and Laughlin 2000, Suarez and Koch 1989). Since the rod-cone retinal model degrades the temporal structure of the stimulus, the Hassenstein-Reichardt correlator “loses” the temporal information it needs to correlate the signals between the two input channels and encode them accurately as motion.

### Visual perception in peripheral retina and natural behavior

We note that the rod and cone kinetic data was fit to data collected from peripheral neurons in primate retina. In many of the psychophysics experiments of motion detection in mesopic conditions, subjects are instructed to discriminate motion that is peripheral to a fixation point in their field of view (Bilino et al., 2008, Yoshimoto and Takeuchi, 2013, Yoshimoto et al., 2016, Sepulveda et al., 2021). One can imagine that in a natural setting, an animal without any behavioral constraints would determine the direction of a moving object by turning to look at the object directly (ie., foveating it). Because our model essentially simulates two nearby ON parasol ganglion cells in the peripheral retina, it is not suited to make predictions about foveal visual processing. Despite this, our model is still well-suited to examine how parallel processing at the lowest level of human vision might impact a more complex behavior. Studies show that subjects perceive the motion of a stimulus as reversing under low light conditions when a blank frame is interspersed in the stimulus, but only when the stimulus is in their periphery (Takeuchi and De Valois, 2009). This is likely due to a first-order biphasic filtering of the stimulus consistent with rod-derived responses (Snowden et al, 1995). Our model ultimately captures a known visual phenomenon, namely mesopic perceptual impairments in peripheral vision, and it demonstrates how interaction between rod- and cone-mediated signal kinetics directly impact motion discrimination.

Specific perceptual deficits in mesopic motion processing have been reported in previous literature with implications for everyday tasks such as driving (Wood, 2019). Motion stimuli that emulate a locomoting figure among a background of incoherent dots were uniquely difficult for human observers to perceive in mesopic conditions, compared to photopic or scotopic conditions (Bilino et al., 2008, Sepulveda et al., 2021). Although it has been posited that rod-cone interactions occurring within the peripheral retina may underlie the phenomenon (Bilino et al., 2008), there has not been a direct examination of responses at the level of the neural circuit. Our model results are in agreement with previous studies which have described perceptual deficits for motion, specifically in dim lighting conditions (Bilino et al., 2008, Yoshimoto et al., 2013, Yoshimoto et al., 2016, Mayeur et al., 2008, Grossman and Blake, 1999, Sepulveda et al., 2021).

## Acknowledgements

Electrophysiological recordings used to train the model were collected by Fred Rieke and William Grimes. Feedback and comments on study from Rieke lab members, including test readers Alison Weber and Alex White. This work was supported by the National Institutes of Health, Bethesda MD, grants K22NS104187, EY028111, and 5R90DA033461, as well as the UW Institute of Neuroengineering Washington Research Foundation Innovation Post-Baccalaureate Fellowship.

## Contributions

FR, GJG, and ASA designed the simulations and the computational study. FR collected the data used to fit the model circuits. ASA created the model. ASA and GJG implemented the simulations, analyzed the simulated data, and drafted the manuscript. All authors participated in editing the manuscript.

---

**Supplemental Figure S1. (Supplement to Fig. 2).**
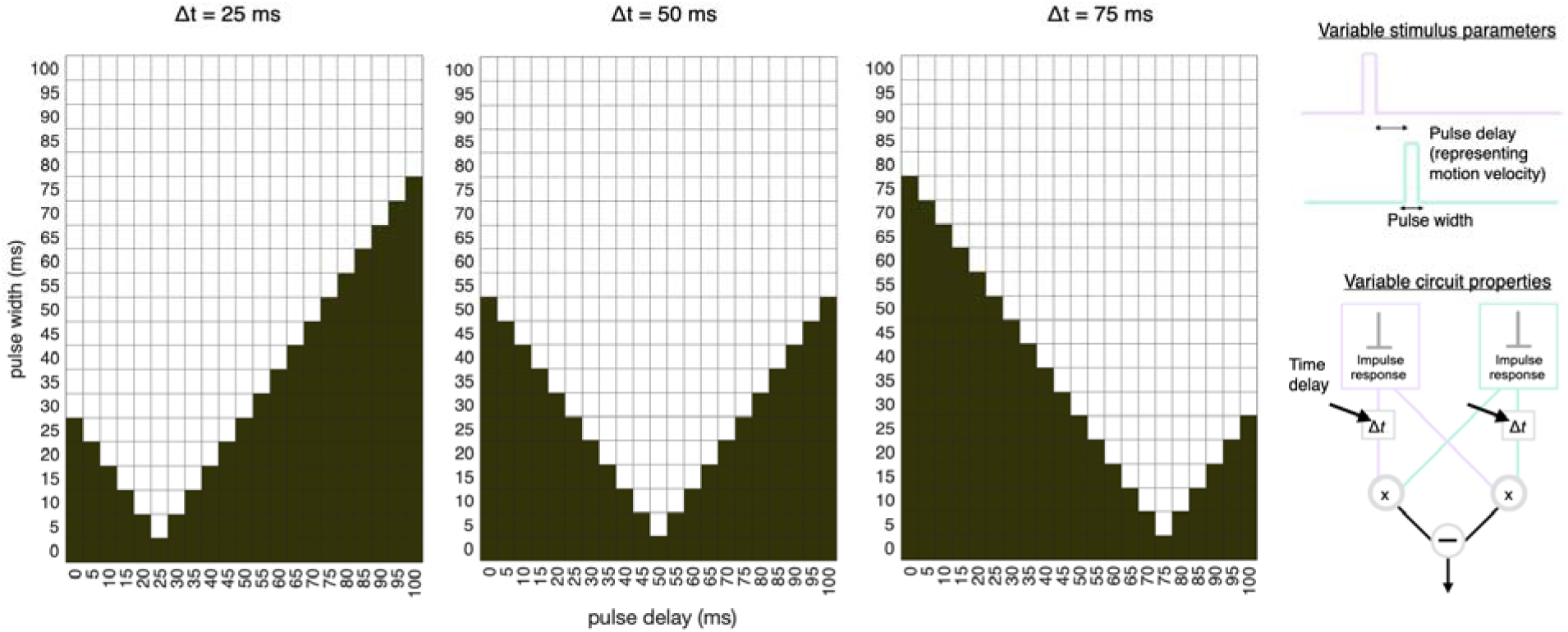
Stimulus correlations in Hassenstein-Reichardt circuit without rod-cone kinetics. We examined whether there was overlap (ie., correlation) for noiseless motion stimuli delivered between the two input channels of a simplified implementation of the correlator circuit, based on the time delay of the circuit and the pulse widths and pulse delays (representing motion velocity) of the stimuli. The dark squares on the grid signify width and delay combinations in which there was no overlap, while the white squares indicate where overlap between the two input channels was possible, and hence correlation could be computed by the circuit. For example, with pulse widths of 0 ms, overlap between the input channels was impossible.

**Supplemental Figure S2. (Supplement to Figure 4).**
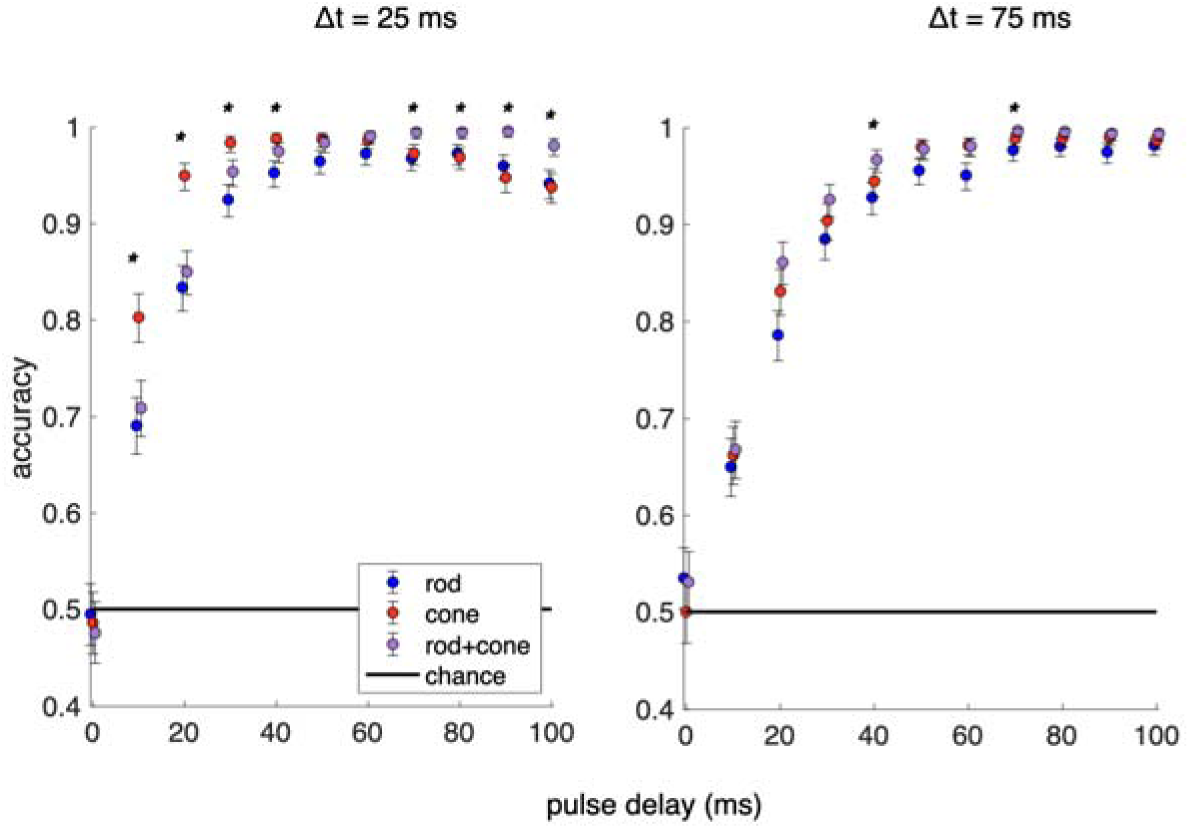
The correlator time delay and velocity tuning. We compare how motion direction discriminability is affected by the Hassenstein-Reichardt correlator time delay (□*t*) in rod, cone, and rod-cone model circuits. We compare two time delays, 25 ms and 75 ms, in contrast to the default time delay we used in all other experiments, 50 ms. (left) With a 25 ms time delay, there is a clear separation between the performances of the cone, rod, and combined rodcone model circuits for stimuli with pulse delays between 10-30 ms. (right) With a 75 ms time delay, there was no statistically significant separation in performance for pulse delays between 10-30 ms. Error bars represent the confidence interval with a minimum coverage of 95% (Clopper-Pearson interval). Solid black line indicates chance (a motion discrimination accuracy of 0.5, ie., 50%). Asterisks denote statistical significance was reached in the comparison between the cone-only circuit performance and the combined rod-cone circuit performance (p ≤ 0.05).

